# Matrilineal Transmission of Familial Excess Longevity (mtFEL): Effects on Cause-specific Mortality in Utah, 1904 - 2002

**DOI:** 10.1101/361881

**Authors:** Elizabeth O’Brien, Richard M. Cawthon, Ken R. Smith, Richard A. Kerber

## Abstract

The heritable component to a long and healthy life is likely to involve the actions and interactions of both nuclear and mitochondrial genetic variants. Using computerized genealogical records with accompanying cause of death information from the Utah population, we previously reported cause-specific mortality rate distributions associated with the nuclear genetic component of familial exceptional longevity. Here we identify Utah matrilineages (mitochondrial lineages) in which overall survival is better than expected, and compare cause-specific mortality rates in those matrilineages to cause-specific mortality rates in the general population. We also examine the effects on cause-specific mortality of interactions between the nuclear and mitochondrial components of familial excess longevity (nuclear *FEL* and *mtFEL*). Among individuals from the bottom quartile of nuclear *FEL*, those who were also in the top quartile for *mtFEL* had lower all-cause, heart disease, cancer, stroke, and diabetes mortality rates than those in the bottom quartile of *mtFEL*. In contrast, among individuals from the top quartile of nuclear *FEL*, the mortality rates from these diseases were similar for those also in the top quartile of *mtFEL* vs. those also in the bottom quartile of *mtFEL*, with the exception of diabetes mortality, which was dramatically suppressed in the high nuclear *FEL* + high *mtFEL* group as compared to the high nuclear *FEL* + low *mtFEL* group. Moreover, the highest mortality rates from diabetes were found in individuals aged 90 years or older who were members of both the high nuclear *FEL* and low *mtFEL* quartiles. These results support the hypothesis that some nuclear genetic variants contributing to long life carry an increased risk of dying from diabetes that is strongly ameliorated by some mitochondrial DNA variants.

## Introduction

Progress in recent years in the development of gene technologies, and success in sequencing the human genome, have been instrumental in advancing our grasp of genes well beyond their sequential dimension. New knowledge that has resulted from these innovations has found many direct applications in areas of biomedical research; and as a growing number of species’ genomes have been sequenced, evolutionary studies have found new momentum, as well. The broad reach of recent progress in gene technologies has naturally forged new crossroads where biomedical and evolutionary studies now meet and look like mutual areas of interest. This study addresses human lifespan, one example at the intersection of biomedical studies of aging and evolutionary studies of life history traits.

Although the evolution of human life history traits has had a long history itself, the thrust has remained largely theoretical due to how long the human lifespan is in relation to how truncated the data usually available to study it. Still the current literature shows how quickly an expanded aperture to the human and other genomes has thrown open the question of how traits, such as lifespan, are genetically regulated and by what specific genetic factors.

Observed variation among species in metabolic characteristics, body size, rates of aging, and maximum lifespan, have suggested experiments in model organisms to identify genes that contribute to the control of these traits. The first experiments in which induced gene mutations were shown to increase lifespan were conducted in worms (Lin, Dorman et al. 1997), flies, and mice; later in mole-rats (Buffenstein and Jarvis 2002; O’Connor, Lee et al. 2002; Woodley and Buffenstein 2002) and fish. Many studies of lifespan heritability in humans have established that up to 25% of the variation is heritable. This portion in humans then, is potentially under genetic control and the genes of interest may well include human homologues of those identified in model systems.

In an earlier paper we analyzed interactions between familial longevity and cause-specific mortality rates in a large Utah population cohort (O’Brien, Kerber et al. 2007). In that study we addressed the hypothesis that genetic contributions to the regulation of human lifespan might do so by influencing rates of aging. Our evidence targeted variation in rates of mortality from the leading age-related causes of it, and we demonstrated that individuals from longer-lived families had lower rates of death from all of ten leading causes, except cancer. Mortality pattern variation was also depicted by age and gender in the population, and the results were consistent with the idea that “longevity genes” might modify lifespan by altering rates of aging through mechanisms that delay the onset or progression of disease.

This study continues our exploration of familial longevity and mortality in the same large Utah population cohort, but here the problem is expressed in terms of mitochondrial longevity. Mitochondria are the sites of energy production and transport. Their role in aging has become the subject of close attention recently as researchers have increasingly understood how this essential function interacts with the regulatory pathways that govern cell life, health, and death. Although the basic chemistry and physics of energy transformation by oxidative phosphorylation were worked out a long time ago, the wider biology of how this essential function contributes to the healthspan of cells, overall somatic condition, and characteristic longevity, is developing in more recent research, both in terms of disease pathology and “normal” aging.

Originating in the oocyte, mitochondria and the multiple small circular genomes (mtDNA) that each contains are maternally transmitted, as are germline mutations in these miniature genomes. Because there are hundreds of mitochondria per cell, each with multiple mtDNA genomes, individual cells host small populations of mitochondrial genomes. New variants can arise during mtDNA copying associated with mitosis, and over cell generations become distributed differentially by selection and random processes, among mitochondria and cells. MtDNA mutations can also occur during mtDNA copying associated with mitogenesis and mitophagy in post-mitotic tissues, and during mtDNA repair following damage from various sources, e.g. by reactive oxygen species (ROS), themselves byproducts of the main energy producing function of mitochondria. Although not without controversy, it’s believed that post-mitotic cells accumulate ROS over time due to declines in electron transport and oxidative phosphorylation efficiency with age (Wallace 2005). Increased ROS can cause the mitochondrial permeability transition pore to open, causing loss of the hydrogen ion gradient across the inner mitochondrial membrane, and triggering apoptosis (programmed cell death), or cellular senescence, depending on the cell type, and, consequently, increasing tissue dysfunction with age, and subsequently the onset of various age-related diseases, such as diabetes, heart disease and stroke. A mechanism of this type might be said to regulate the rate at which an organism ages, the rate of age-related disease onset or progression, and ultimately, rates of death.

In this study we further test the dynamic between familial longevity and familial mortality with respect to cause-specific mortality risks. As in other populations with a similarly western lifestyle and medical practices, the highest incident causes of mortality in Utah are three aging-related diseases -- heart disease, cancer, and stroke, while diabetes is the fifth highest incident cause of death (after influenza/pneumonia). In this study we look again at these most common causes of death, and we measure individual longevity based on one’s family history of longevity. We have previously called this measure “familial excess longevity” (***FEL***), but here we modify it to express longevity through matrilineal descent alone. We call this adaptation, “***mtFEL***”.

## Methods

### Utah Population Database (UPDB) and cause of death data

Data for this study were drawn from the UPDB population resource at the University of Utah. The collection history and data inventory of the UPDB is described in detail elsewhere (Skolnick 1980; Jaro 1989). Source data used for vital status follow-up information includes a variety of data linked to the UPDB: state driver license; Centers for Medicaid and Medicare Studies (CMS); Social Security Death Index (SSDI); and Cancer Registries for the states of Utah and Idaho. Our use of certified death records for the state of Utah for coded causes of death for this analysis is also described fully elsewhere (O’Brien, Kerber et al. 2007; Kerber, O’Brien et al. 2008).

### Study cohort

A cohort was selected to include all persons born 1830-1963, who have at least one family member in the database. Each member of the cohort lived or died in Utah since 1904, when standardized state death certification was first used. Using the data sources listed above, we confirmed the vital status to age 40 of every cohort member. Defined this way, the cohort included more than 766,000 members followed through 2003.

The extensive family histories available in UPDB allow us to address familial trait clustering in the population only with efficient means of capturing information while navigating these very deep and intertwined pedigrees. We departed from the tradition of using minimal pedigree data for genetic and epidemiologic studies some years ago (O’Brien, Kerber et al. 1994; Kerber 1995). Instead we have developed methods to measure the familiality of a trait using full relationship networks over the deep pedigrees of the UPDB (Kerber, O’Brien et al. 2001; Kerber and O’Brien 2005; O’Brien, Kerber et al. 2007).

### Familial standardized mortality ratio (FSMR)

Familial standardized mortality ratios were calculated for all individuals in the cohort, similar to the Familial Standardized Incidence Ratio, or FSIR (Kerber 1995). FSMR is the observed mortality among family members of an individual, divided by the expected mortality, with weights for relatives *j* of a proband *i* given by the kinship coefficient (Malecot 1948), the probability that *i* and *j* share a given gene identical by descent from a common ancestor, *f(i,j)*:

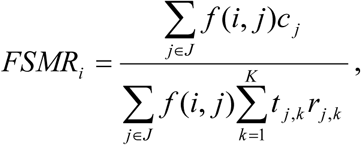

where *J* is the set of all relatives of *i*, *c_j_* is an indicator variable (1 if *j* dies of the disease of interest, 0 otherwise), *t_j,k_* is the time spent by *j* in the *k*^th^ risk stratum (designated by sex and age), and *r_j,k_* is the population risk per unit time for a person in stratum *k*. FSMR measures an individual’s family history of a trait, and the approach has been shown elsewhere to be a useful predictor of disease risk (Slattery and Kerber 1993; Slattery and Kerber 1994; Kerber and O’Brien 2005). Here it’s used to measure the familial component of cause-specific mortality.

### Familial excess longevity (FEL)

Familial excess longevity was originally presented as a means of quantifying individual excess longevity based on cohort-adjusted expectations, as well as a family history of longevity (Kerber, O’Brien, Cawthon, unpublished, ca. 1998). A complete description can be found elsewhere (Kerber, O’Brien et al. 2001), and in more recent studies of age-related mortality outcomes (Garibotti, Smith et al. 2006; O’Brien, Kerber et al. 2007; Kerber, O’Brien et al. 2008). First, *excess longevity (EL)* was estimated for all individuals in the cohort, defined as the difference between an individual’s attained age and the age to which that individual was expected to live incorporating into the model potential confounders of longevity. Here we include gender and birth year, although other environmental or behavioral factors that influence lifespan could be included as well. The expected longevity (*ŷ*) is estimated from an accelerated failure time model in the following manner:

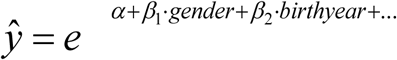

where *α* is the intercept, *β_1_*…*β_n_* are slope coefficients, and the excess longevity (*l*) is simply *y* − *ŷ*, with *y* the attained age in years assessed for birth cohorts (e.g., born <1900) for which exceptional longevity is possible. This is similar to the method of Bocquet-Appel to estimate the heritability of longevity at Arthez d’Asson (Bocquet-Appel 1990).

Next, *familial excess longevity* (Kerber, O’Brien et al. 2001) is estimated for each individual in the cohort. An individual’s *FEL* is a summary calculation of excess longevity over all of an individual’s family members who lived to the baseline age of 65. At age 65 and over death rates are increasingly age-related, and more specifically reflect variation in senescence apart from causes of early mortality. We average excess longevity of an individual’s family members, weighted by kinship coefficients, to yield an estimate of *FEL*:

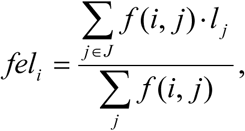

where *fel_i_* is the excess longevity for subject *i*, *J* is the set of all relatives of *i, l_j_* is the excess longevity of the *j*th member of *J*, and *f(i,j)*, the probability that *i* and *j* share a gene identical by descent from a common ancestor.

Matrilineal FEL (*mtFEL*) is a simple variant of *FEL*: it is the mean *EL* across all strictly matrilineal relatives, weighted equally. This computation reflects the inheritance pattern of mitochondrial DNA. We can estimate the independent effects of nuclear and mitochondrial inheritance on mortality by fitting a proportional hazard model that incorporates both *FEL* and *mtFEL*. In principle, both measures can be confounded by shared environment, although *FEL* is more susceptible to such confounding because close relatives are more influential in the computation of *FEL* than in the computation of *mtFEL*.

We use proportional hazards models to estimate the overall effects of variation in longevity among relatives (matrilineal or otherwise) on mortality. We are also interested in possible interactions between nuclear and mitochondrial variants, and in the age distribution of effects of variation in the nuclear and mitochondrial genomes. To test for interaction between nuclear and mitochondrial variants, we fit proportional hazards models using the contrasting tails of the *FEL* and *mtFEL* distributions. We defined four subgroups of cohort members: 1) those in the bottom quartile for both *FEL* and *mtFEL*; 2) those in the highest quartile of *FEL* but the lowest quartile of *mtFEL*; 3) those in the lowest quartile of *FEL* but the highest quartile of *mtFEL*; and 4) those in the highest quartile for both *FEL* and *mtFEL*. All proportional hazards models included terms for sex (female vs. male), year of birth, and *FSMRs* for heart disease, cancer, stroke, and diabetes, defined as described above, as well as *FEL* and *mtFEL*.

We also fit generalized gamma survival regression models for each of these four groups, in order to examine whether the effects of *FEL* and/or *mtFEL* vary with time. Generalized gamma models are described in more detail in (Garibotti, Smith et al. 2006; O’Brien, Kerber et al. 2007; Kerber, O’Brien et al. 2008); we use them here to estimate cause-specific mortality hazard functions without assuming that the effects of covariates are constant over time.

## Results

Table 1 shows estimates of relative risk, in the form of hazard rate ratios (HRR), for death from all causes and four of the leading causes of death in the UPDB cohort (heart disease, cancer, stroke, and diabetes). Risk estimates are given for sex (females as compared to males); year of birth; *FSMR* for heart disease, cancer, stroke, and diabetes; and *FEL* and *mtFEL*. For all terms except sex, the HRR estimate given in Table 1 is for a change of +1 standard deviations, designed to reflect a typical, rather than an extreme, difference between individuals. The 95% confidence interval excludes 1.0 for almost every effect estimate in Table 1. The exceptions are: *FSMR Heart* (All Causes), *FSMR Diabetes* and *mtFEL* (Cancer), and *mtFEL* (Stroke). For all causes of death except cancer, a change of one standard deviation in *FEL* (about 2.4 years) results in a drop of about 10% in risk of death, after adjusting for family history for each of the four leading causes of death and *mtFEL*. The corresponding change in *mtFEL* results in a smaller and more variable reduction in risk of death: 3% overall, 4-5% for heart disease and diabetes, and no observable reduction in risk for cancer or stroke.

**Table 1.** Hazard rate ratios (HRR) and confidence intervals for all cause mortality, heart disease, cancer, and diabetes associated with sex, year of birth, and a one standard deviation change in family history of each disease (FSMR), familial excess longevity (FEL), or matrilineal familial excess longevity (mtFEL).

| Effect | All Causes |  | Heart Disease |  | Cancer |  | Stroke |  | Diabetes |  |
| --- | --- | --- | --- | --- | --- | --- | --- | --- | --- | --- |
|  | HRR | 95% CI | HRR | 95% CI | HRR | 95% CI | HRR | 95% CI | HRR | 95% CI |
| Sex (female) | 0.72 | (0.71-0.72) | 0.65 | (0.64-0.66) | 0.83 | (0.81-0.84) | 1.02 | (1.00-1.05) | 1.17 | (1.11-1.23) |
| Year of birth | 0.75 | (0.75-0.76) | 0.68 | (0.67-0.68) | 1.11 | (1.09-1.12) | 0.64 | (0.63-0.65) | 1.12 | (1.08-1.16) |
| FSMR Heart | 1.00 | (0.99-1.00) | 1.23 | (1.22-1.24) | 1.03 | (1.02-1.04) | 1.17 | (1.15-1.18) | 1.09 | (1.05-1.12) |
| FSMR Cancer | 0.96 | (0.96-0.96) | 1.03 | (1.02-1.04) | 1.18 | (1.16-1.19) | 1.03 | (1.01-1.04) | 1.04 | (1.01-1.08) |
| FSMR Stroke | 0.97 | (0.97-0.98) | 1.05 | (1.03-1.05) | 0.98 | (0.97-0.99) | 1.13 | (1.11-1.14) | 1.04 | (1.01-1.06) |
| FSMR Diabetes | 1.01 | (1.00-1.01) | 1.04 | (1.03-1.05) | 0.99 | (0.98-1.01) | 1.04 | (1.03-1.06) | 1.41 | (1.38-1.44) |
| FEL | 0.89 | (0.88-0.89) | 0.90 | (0.90-0.91) | 0.98 | (0.96-0.99) | 0.91 | (0.90-0.93) | 0.90 | (0.88-0.93) |
| mtFEL | 0.97 | (0.97-0.98) | 0.96 | (0.95-0.97) | 0.99 | (0.98-1.01) | 0.99 | (0.97-1.00) | 0.95 | (0.92-0.98) |

Table 2 gives relative risk estimates for the tails of the joint distribution of *FEL* and *mtFEL*. All comparisons are relative to the cohort members in the lowest quartiles of both *FEL* and *mtFEL*. This analysis shows more clearly the effects of variation in *mtFEL* against backgrounds of high or low *FEL* and vice versa. In all cases, the effect of high *FEL* alone is larger than the effect of high *mtFEL* alone. For all cause mortality, heart disease, and cancer, those in the highest quartile of *mtFEL* but the lowest quartile of *FEL* are at significantly reduced risk of death compared to those in the lowest quartile of both. For stroke and diabetes, there is no significant difference between the highest and lowest quartiles of *mtFEL* among those in the lowest quartile of *FEL*. Among those in the highest quartile of *FEL*, however, the case is quite different. The risk of death from diabetes is substantially reduced among those in the highest quartile of both *FEL* and *mtFEL*, as compared to those in the highest quartile of *FEL* but the lowest quartile of *mtFEL*. There is also a small but significant additional reduction in risk of all cause mortality between the high *FEL*-low *mtFEL* and high *FEL*-high *mtFEL* groups.

**Table 2.** Hazard rate ratios (HRR) and confidence intervals for all cause mortality, heart disease, cancer, and diabetes associated with membership in the lowest or highest quartile of FEL and mtFEL, adjusted for the risk factors in Table 1.

| Effect | All Causes |  | Heart Disease |  | Cancer |  | Stroke |  | Diabetes |  |
| --- | --- | --- | --- | --- | --- | --- | --- | --- | --- | --- |
|  | HRR | 95% CI | HRR | 95% CI | HRR | 95% CI | HRR | 95% CI | HRR | 95% CI |
| Low FEL/Low mtFEL | 1.00 |  | 1.00 |  | 1.00 |  | 1.00 |  | 1.00 |  |
| Low FEL/High mtFEL | 0.91 | (0.89-0.93) | 0.86 | (0.82-0.90) | 0.92 | (0.85-0.99) | 0.95 | (0.87-1.04) | 0.94 | (0.80-1.11) |
| High FEL/Low mtFEL | 0.67 | (0.65-0.68) | 0.64 | (0.61-0.67) | 0.90 | (0.84-0.96) | 0.74 | (0.67-0.81) | 0.76 | (0.64-0.90) |
| High FEL/High mtFEL | 0.64 | (0.63-0.64) | 0.63 | (0.62-0.65) | 0.92 | (0.88-0.96) | 0.71 | (0.68-0.75) | 0.58 | (0.52-0.64) |

Figures 1-5 show generalized gamma estimates of the cause-specific mortality hazard function for the four risk groups and five cause of death categories given in Table 2.

**Figure 1.**
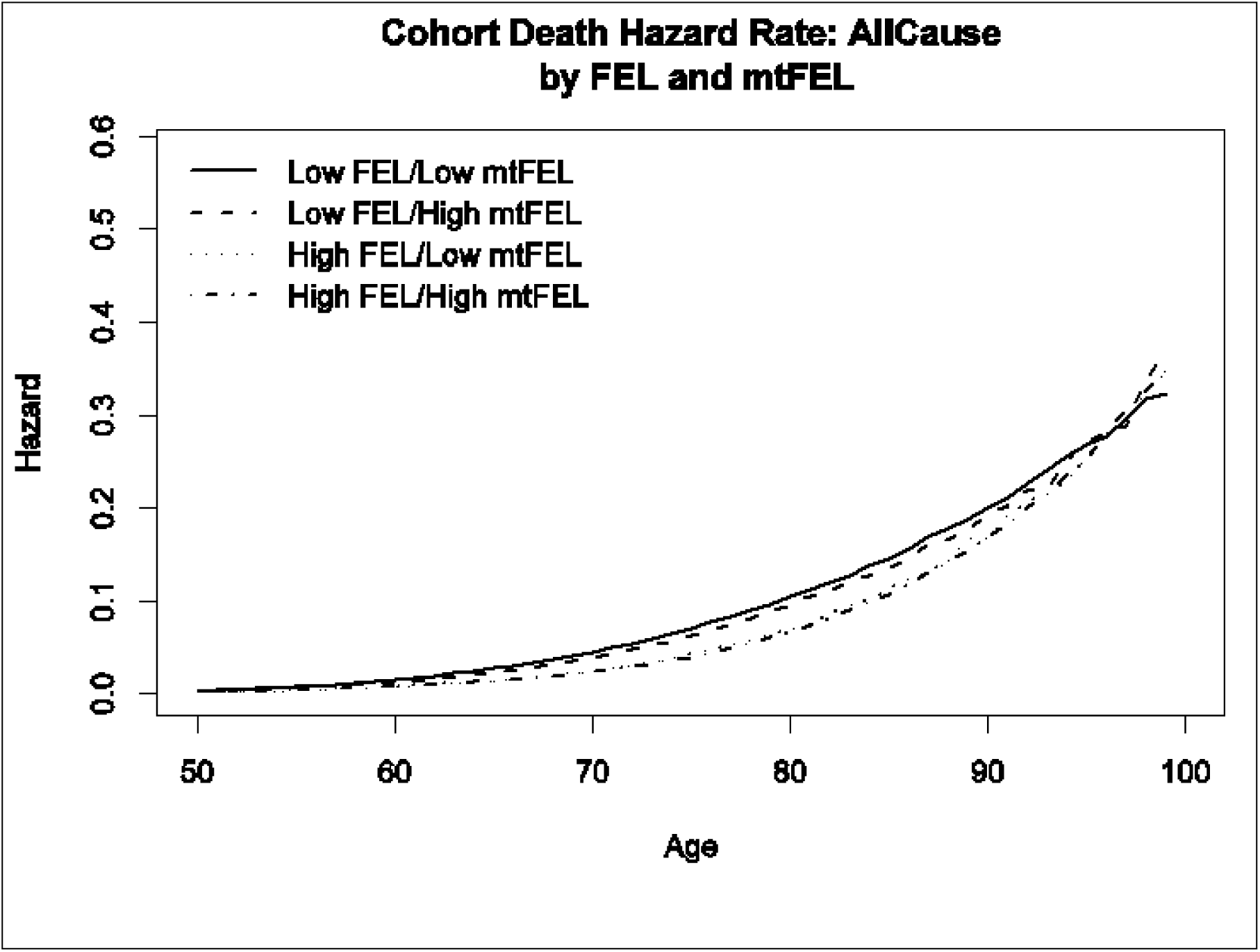

**Figure 2.**
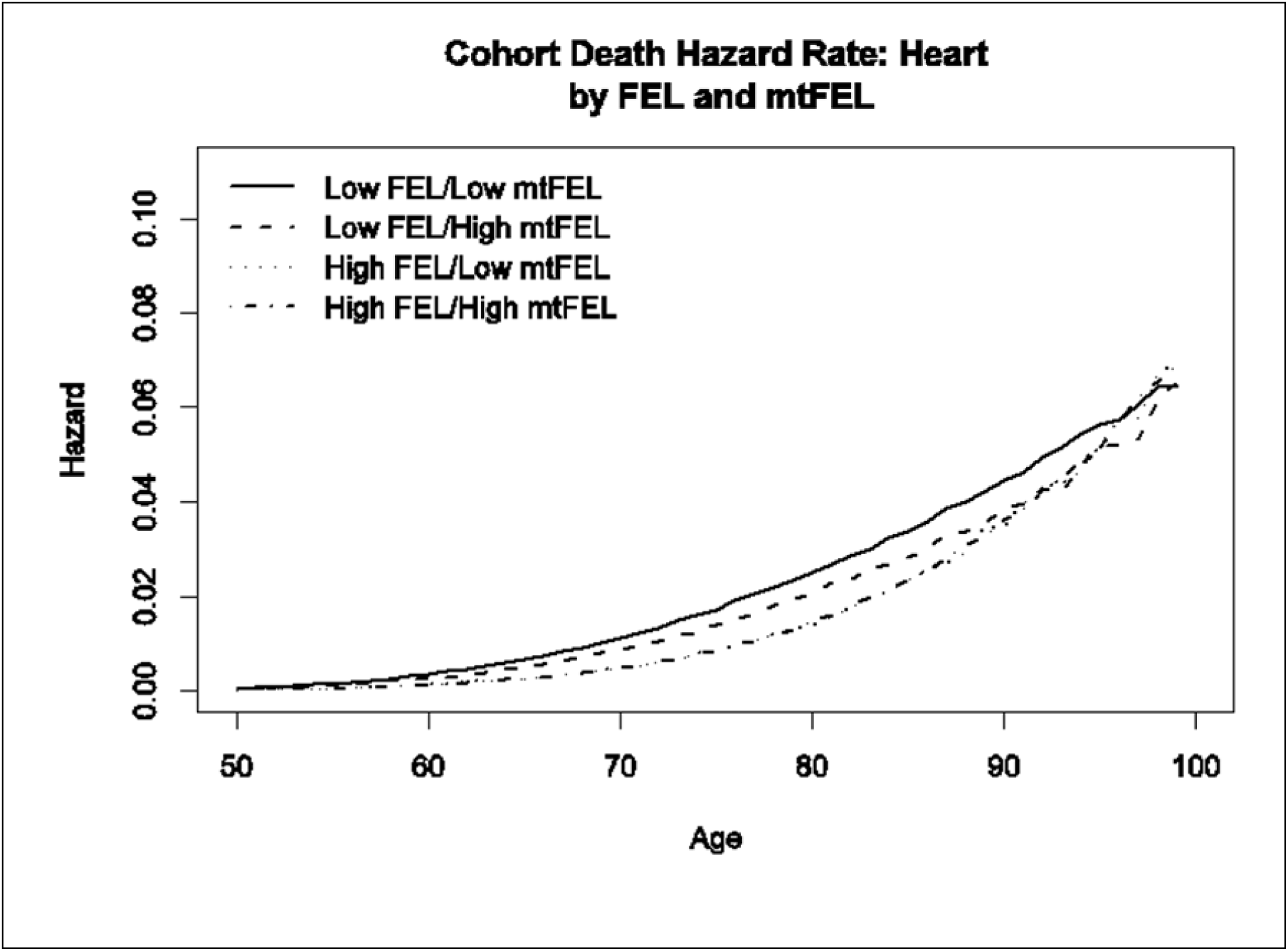

**Figure 3.**
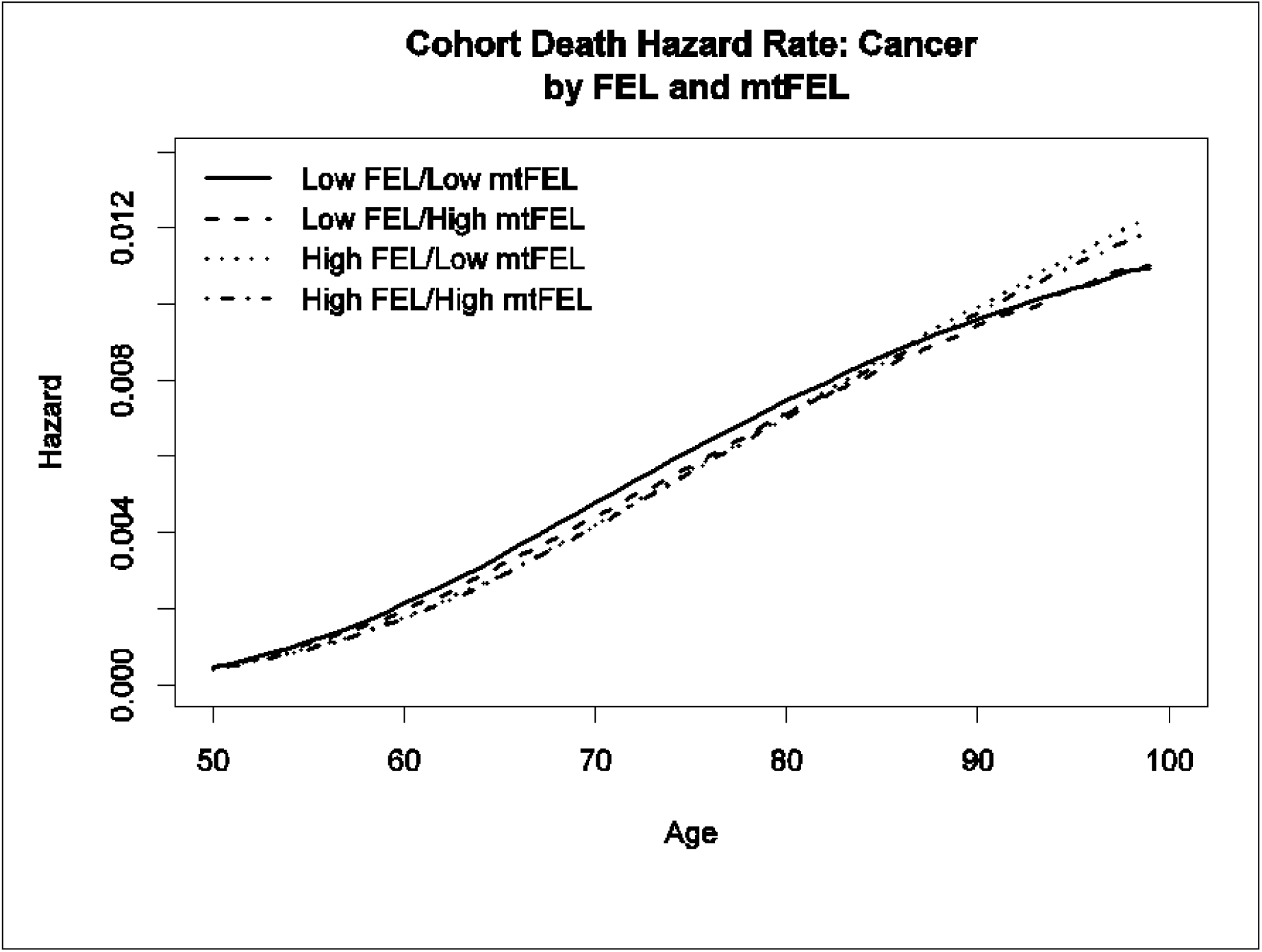

**Figure 4.**
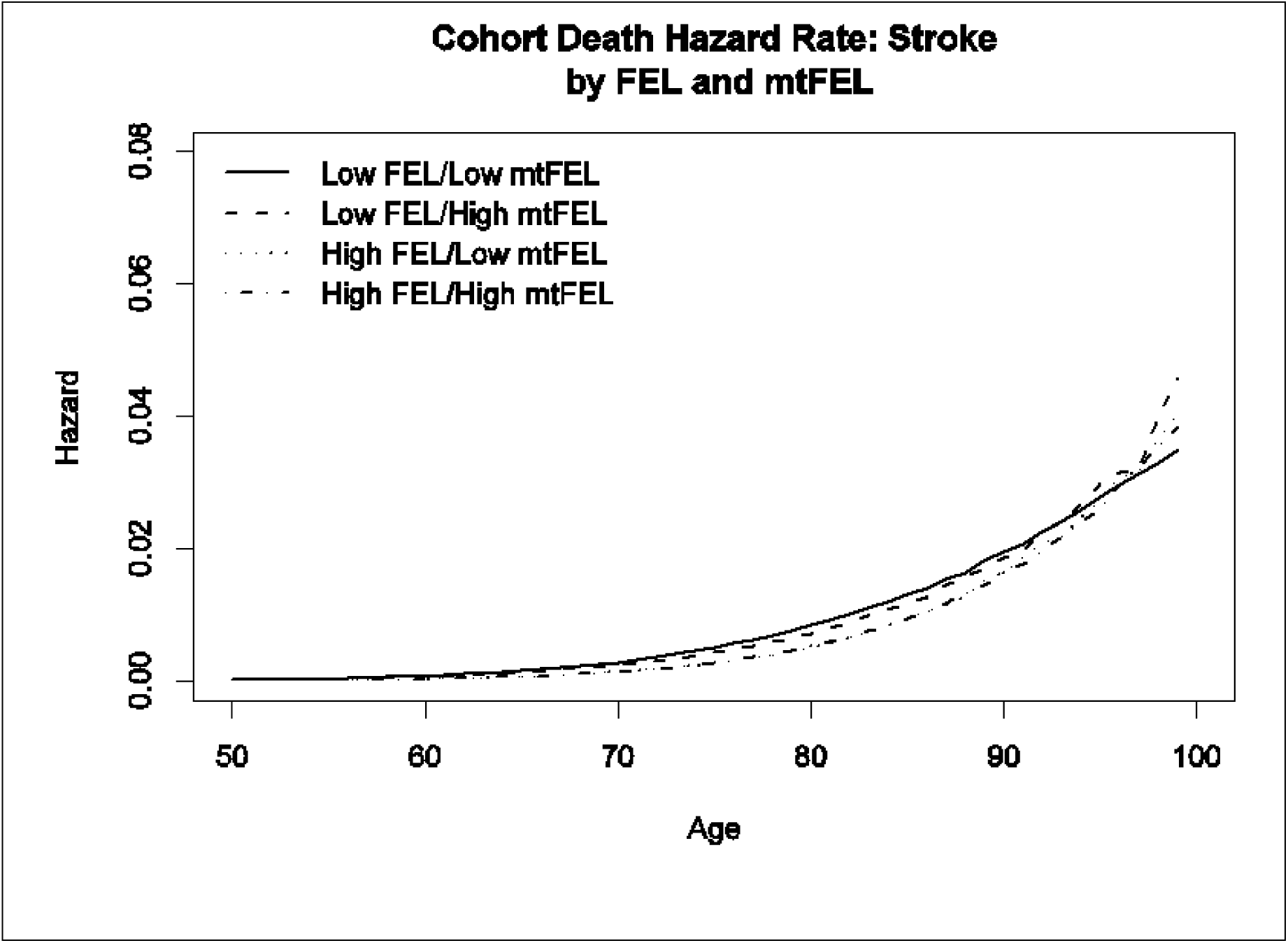

**Figure 5.**
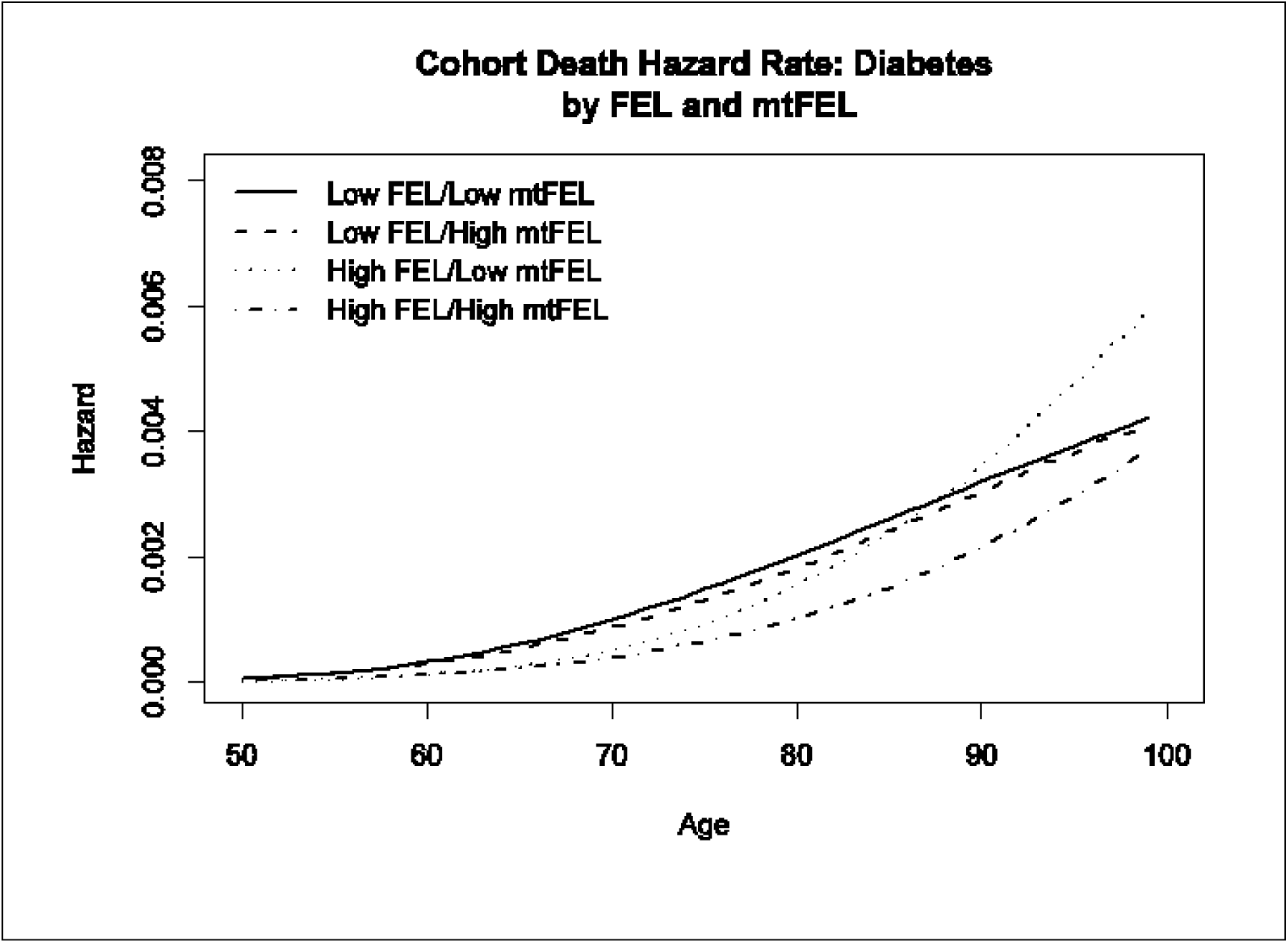

## Discussion

Insulin and insulin-like growth factor signaling pathways regulate aging and lifespan across multiple species (worms, flies, and mammals). The daf-2 gene in the nematode *C. elegans* is homologous to the human genes encoding the insulin receptor and insulin-like growth factor I receptor (Kimura et al. 1997). Some amino acid changing mutations in the daf-2 gene dampen signaling through its encoded receptor, resulting in a decrease in the inhibition of daf-16, which when active contributes to increased lifespan via several mechanisms (Lapierre and Hansen 2012). Some amino acid changing mutations in the insulin receptor gene in human patients with insulin resistance correspond directly to the lifespan-extending mutations in daf-2 (Kim et al. 1992; Kimura et al. 1997). Centenarians, as compared to younger controls, are more frequent carriers of amino acid changing mutations in the insulin-like growth factor I receptor gene that decrease IGFI signaling (Suh et al. 2008). Humans have four genes homologous to daf-16: FOXO1, FOXO3, FOXO4, and FOXO6. Genetic variants in FOXO3 have been shown to be increased in frequency in independent cohorts of human centenarians, as compared to controls (Willcox et al. 2008; Flachsbart et al. 2009). Finally, activated FOXO genes, when combined with a high fat diet, contribute to the development of insulin resistance and diabetes in mice (Nakae et al. 2002; Kim et al. 2009; Nwadozi et al. 2016).

Therefore, it is reasonable to hypothesize that some human nuclear genetic variants that decrease insulin/IGF signaling and/or activate one or more of the FOXO genes directly will either increase the healthspan and lifespan, or increase the risk of developing Type 2 Diabetes, or both. The life course followed by carriers of such conditionally beneficial nuclear genes may strongly depend on life style factors, such as caloric intake and exercise, but also may depend on which mitochondrial DNA sequence variants have been inherited. Further studies to investigate differences in the gene expressions, biochemistry, and metabolism of individuals with high nuclear *FEL* + high *mtFEL* vs. individuals with high nuclear *FEL* + low *mtFEL* may add significantly to our understanding of the molecular mechanisms of aging, and lead to the development of medical interventions to increase the human healthspan and lifespan.

## Acknowledgements

We thank the Pedigree and Population Resource (funded by the Huntsman Cancer Foundation) for its valuable role in the ongoing collection, maintenance, and support of the Utah Population Database.

